# *In vivo* optogenetics using a Utah Optrode Array with enhanced light output and spatial selectivity

**DOI:** 10.1101/2024.03.18.585479

**Authors:** Niall McAlinden, Christopher F. Reiche, Andrew M. Clark, Robert Scharf, Yunzhou Cheng, Rohit Sharma, Loren Rieth, Martin D. Dawson, Alessandra Angelucci, Keith Mathieson, Steve Blair

## Abstract

Optogenetics allows manipulation of neural circuits *in vivo* with high spatial and temporal precision. However, combining this precision with control over a significant portion of the brain is technologically challenging (especially in larger animal models). Here, we have developed, optimised, and tested in vivo, the Utah Optrode Array (UOA), an electrically addressable array of optical needles and interstitial sites illuminated by 181 µLEDs and used to optogenetically stimulate the brain. The device is specifically designed for non-human primate studies. Thinning the combined µLED and needle backplane of the device from 300 µm to 230 µm improved the efficiency of light delivery to tissue by 80%, allowing lower µLED drive currents, which improved power management and thermal performance. The spatial selectivity of each site was also improved by integrating an optical interposer to reduce stray light emission. These improvements were achieved using an innovative fabrication method to create an anodically bonded glass/silicon substrate with through-silicon vias etched, forming an optical interposer. Optical modelling was used to demonstrate that the tip structure of the device had a major influence on the illumination pattern. The thermal performance was evaluated through a combination of modelling and experiment, in order to ensure that cortical tissue temperatures did not rise by more than 1°C. The device was tested *in vivo* in the visual cortex of macaque expressing ChR2-tdTomato in cortical neurons. It was shown that the strongest optogenetic response occurred in the region surrounding the needle tips, and that the extent of the optogenetic response matched the predicted illumination profile based on optical modelling – demonstrating the improved spatial selectivity resulting from the optical interposer approach. Furthermore, different needle illumination sites generated different patterns of low-frequency potential (LFP) activity.

## 1. Introduction

Optogenetics has made transformational contributions to neuroscience, enabling experiments that can dissect the roles of specific components in neural circuits [1]. Recently, the technique has been translated to clinical trials, where it was used to partially restore light perception in blind patients [2]. The technique relies on several disciplines, including protein and genetic engineering for the development and delivery of light-sensitive ion channels, ion pumps and other neural modulators[3]. Photonic engineering also plays an important role in the development of methods to deliver light to neural substrates through highly scattering tissue with the required spatial and temporal resolution[4].

Advances in optogenetics have progressed rapidly, predominantly focusing on the mouse model. However, progress has not been as rapid in models that lack the sophisticated genetic tools and constructs developed for mice, such as the non-human primate [5–8]. Extending optogenetic methods to primate models is important to increase our understanding of neural functions in brains more similar to humans, and as an essential large animal testbed towards further clinical translation. In addition to the development of opsin delivery toolkits, light delivery methods used in mouse models are not always adequate for the larger primate brain, where neural circuits require optical targeting spread over an increased volume [9, 10]. For experiments where only a single light source is required, current tools consisting of single optical cannula coupled to an LED or laser are sufficient, as long as care is taken to not exceed phototoxic levels[11]. However, delivery of patterned or multi-site illumination is more challenging. µLED probes[12–15] and waveguide devices [16, 17] that work well in mouse studies do not provide light over a broad enough spatial extent for studies in larger mammals[8].

Surface illumination strategies [18–22] have been shown to work in primates, but light penetration depths are limited to less than 1 mm due to scattering and absorption in brain tissue [19]. For studies requiring deeper illumination, important in larger animal models as cortical thickness are typically in excess of 1 mm, several groups have developed penetrating microneedle devices [23, 24]. However, without an integrated light source, the animal must remain head fixed so that optical alignment can be maintained[25]. Other researchers have demonstrated that it is possible to insert multiple fibre optics [26] to achieve patterned light for deep structures, but again maintaining alignment of an external light source is required.

Previously, we reported on the Utah Optrode Array (UOA) [27], a light delivery device that addresses these issues. It consists of a matrix of µLEDs, which is coupled to an array of microneedles, creating a device that can provide optogenetic illumination from 181 individual sites both to deeper layers (needles sites) and the surface of the cortex (interstitial sites). As the µLED source and penetrating needles are directly coupled, optical alignment is maintained by the fabrication process. However, inefficiencies in optical coupling remain a significant challenge, due primarily to the Lambertian nature of µLED emission, where optical efficiencies (µLED to target site) can be as low as 0.2%. This means that the µLED needs to be driven with drive currents of 10-100 mA (to deliver enough light to exceed typical optogenetic activation thresholds), which in turn means short pulse widths or low duty cycles are required to ensure the device remains within safe thermal limits. The Lambertian emission profile also means that optical crosstalk between stimulation sites can be problematic, creating ambiguities in the exact volume of tissue activated and reducing the applicability of the device to neuroscience experiments. Improving this coupling is central to enabling an optoelectronic, multi-site device that can provide spatially discrete optogenetic activation and operate at a high dynamic range (irradiance and duty cycle) without heating the brain by more than 1°C[28, 29].

In this work, a glass needle array is integrated with a thin 80 µm Si interposer, which is directly coupled to a µLED array. The closer proximity of the μLED to the glass needle aperture improves the optical efficiency of light delivery to tissue, allowing the device to be operated at significantly lower drive currents, improving the thermal characteristics and light output properties. For example, this approach produces the same optical output power from the needle tips (80 µW) at half the electrical current of the previous approach (10 mA instead of 20 mA). The optical interposer also reduced stray light from the base of the needles for these drive currents (from 85 µW to 7.2 µW), allowing single-site activation with reduced cross-talk between epicortical and deep sites.

Thermal management to protect neural tissue from heating was further improved by the inclusion of a Dura-Gel (silicone) layer between the active element of the device and the delicate cortical surface. For example, when 10 µLEDs were operated to give an optical output power of 660 µW (sufficient to optogenetically excite neurons expressing ChR2 in a tissue volume of 0.11 mm^3^) the previous approach had a thermally imposed duty cycle limit of 7%, which is extended to 25% with this new generation device. These increases in the duty cycle and device performance greatly expand the scope of *in vivo* studies that can be pursued [30]. Here we focus on the technical aspects and performance of the device but also provide further demonstration of its functionality with *in vivo* tests in the non-human primate (macaca fascuularis) cortex. Spatially distinct, multi-unit activity driven by different µLED sites is demonstrated.

## 2. Methods

### 2.1 Optical Modelling

To optimise the device design and fabrication, an optical model was created. This model was also used to understand light spread in tissue. Optical ray-tracing software (Zemax-Optics Studio 12, non-sequential mode) was used. Brain tissue was modelled using a Henyey-Greenstein scattering model, with a scattering coefficient of 10 mm^-1^, an absorption coefficient of 0.07 mm^-1^, and an anisotropy of 0.88[31, 32]. To generate the cross-section images from simultaneously illuminated µLEDs, the light output from a single needle was duplicated, spatially translated, and then summed with the output from other needles. The peak optical power out of each needle was then measured experimentally, for a given current/voltage, and used as an input to the model. This allowed for experimental variations in device optical output to be accommodated within the simulated results. To confirm that the optical modelling accurately reproduced experimental results, the optical output was measured by imaging the emission profile in fluorescein and compared with modelled output.

### 2.2 Device Fabrication – Utah Optrode Array

The implantable optogenetic device described here consists of two components: a glass needle array and a µLED array. The optrode array and the µLED array are fabricated separately and integrated during a final device assembly step. The completed device with a single µLED illuminated is shown in Figure 1A.

**Figure 1:**
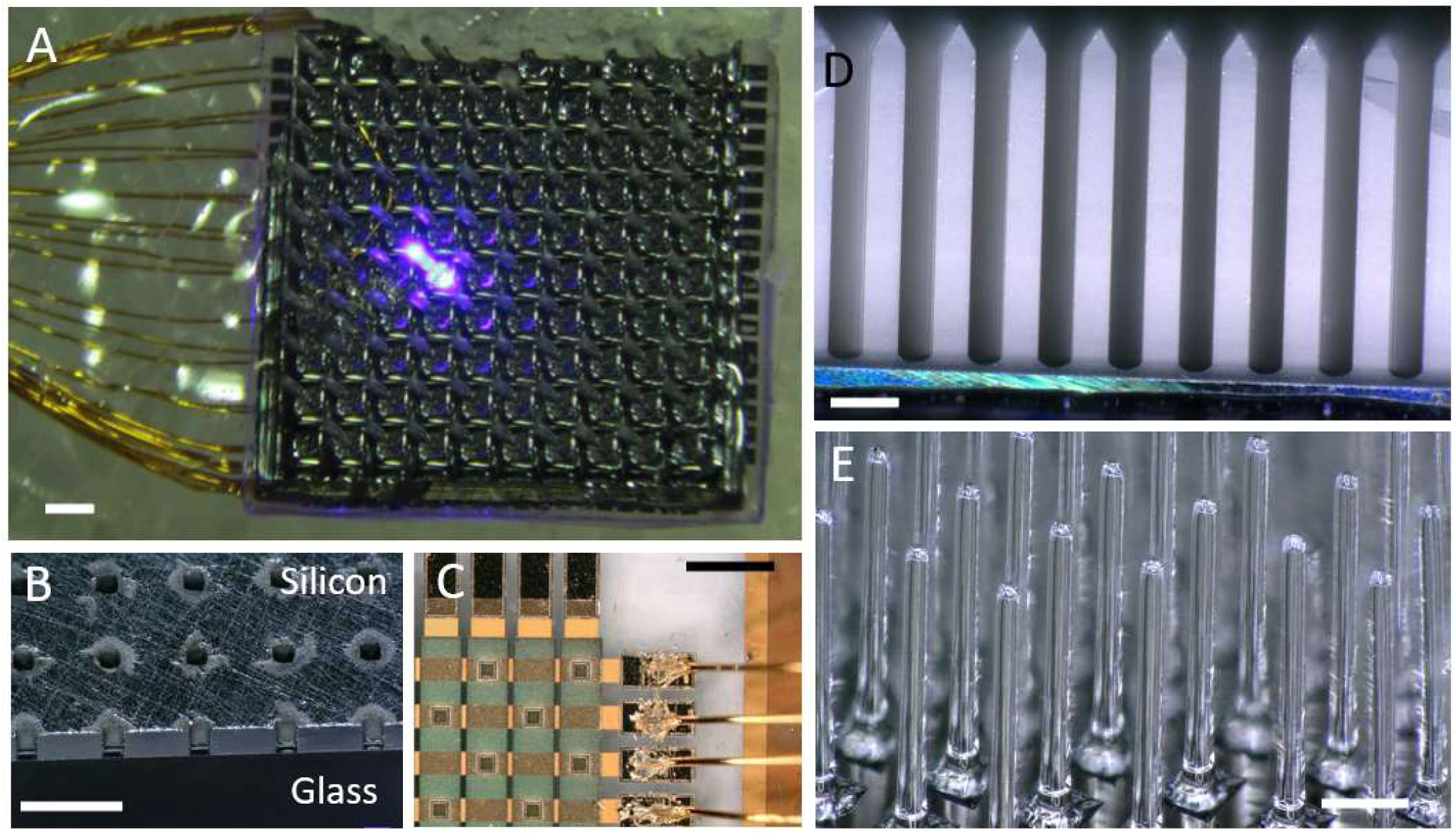
**A** Image of the completed device with a single µLED illuminated (4 mm device width). **B** Cross-section image of silicon interposer with through-silicon vias (TSVs), anodically bonded to the glass optrode base. In this example, TSVs were not opened for interstitial sites. **C** Image of a portion of the µLED device. Wire bonds are visible on the right-hand side of the image. **D** Optrodes after dicing to reveal the needle shape. **E** Finished Utah Optrode Array. The glass needles were chemically etched to reduce their diameter further and thermally threated to smoothen their surface. All scale bars are 400 µm.

The fabrication of the UOA is illustrated in Figure 1. A 100 µm thick silicon wafer is anodically bonded to a 2 mm-thick borosilicate glass wafer (Figure 1B). The Si wafer will form the optical interposer, with the glass wafer being processed into the optical needle array. The anodic bonding was achieved at 350°C, in a low vacuum (10^-3^ mbar), with a force of 2000 N, and a potential difference across the sample of 1000 V for 20 minutes using an EVG 520IS, (EVG, Austria). A thick photoresist (AZ9260, MicroChemicals) was spin-coated on the silicon and standard photolithographic techniques were used to pattern arrays of 80 µm diameter holes forming the mask for through-silicon via (TSV) etching. The TSVs were etched using the Bosch deep reactive ion etching (DRIE) process in a 100 ICP, Oxford Instr. Figure 1B shows the fully etched interposer array. After this step the glass wafer – interposer combination is typically diced and subdivided in smaller dies for the subsequent needle dicing steps.

The UOA fabrication is adapted from previous processes [23] and starts with the glass/silicon interposer wafer. A bevelled dicing saw blade is used to cut the pyramidal-shaped optrode tips into the glass. By using blades with different bevel angles the pyramid angle can be adjusted to produce the desired tip shape and hence light emission profile. The optrode needles are then defined using a straight dicing blade performing cuts with a pitch of 400 µm, an edge width of 200 µm and a height of approximately 1.6 mm, leaving a thin glass backplane (which is removed in the subsequent etch step). This process can be used to create optrode arrays with a range of dimension, in this case 10×10 arrays were produced. Using thicker blades, it is in principle possible to create optrode needles with even smaller edge widths (thinner). However, processing constraints mean that thinner optrodes are more likely to break during the dicing process. Figure 1D shows the needles after dicing on the interposer device, note the optically scattering sidewalls.

To prevent the interposer holes from filling with etchant during the next steps, the die is then mounted onto a carrier wafer using WaferGrip (Dynatex International). A 9:1 mix of hydrofluoric acid (concentration of 49%) and hydrochloric acid (concentration of 37%) is used to thin the optrodes to their target width. This gives a well-controlled process with an etch rate of approximately 6 μm/min at room temperature. After etching, the devices are rinsed with deionized water, removed from the carrier wafer, and cleaned from WaferGrip residues using successive baths of heated xylenes (120 °C), n-butyl acetate (NBA), isopropanol (IPA) and deionized water.

Finally, the cleaned array batch is subjected to thermal treatment under vacuum in a muffle furnace. The device was heated to 560 °C and held for 1 hr to remove stress from the dicing steps. It is then heated to 725 °C and held for 2 hrs which causes some glass surface reflow and gives an optically smooth surface to the needles (Figure 1E). During this step, there is a noticeable geometry change including a slight rounding of the corners of the optrodes and a reduction of their length. Extended annealing causes further rounding of the tips which could be used to change the light emission properties of the device (Figure 2F and G). After this step, the device is again held at 560 °C for 1hr to relieve stress in the glass that forms during cooling and then left to cool to room temperature.

**Figure 2:**
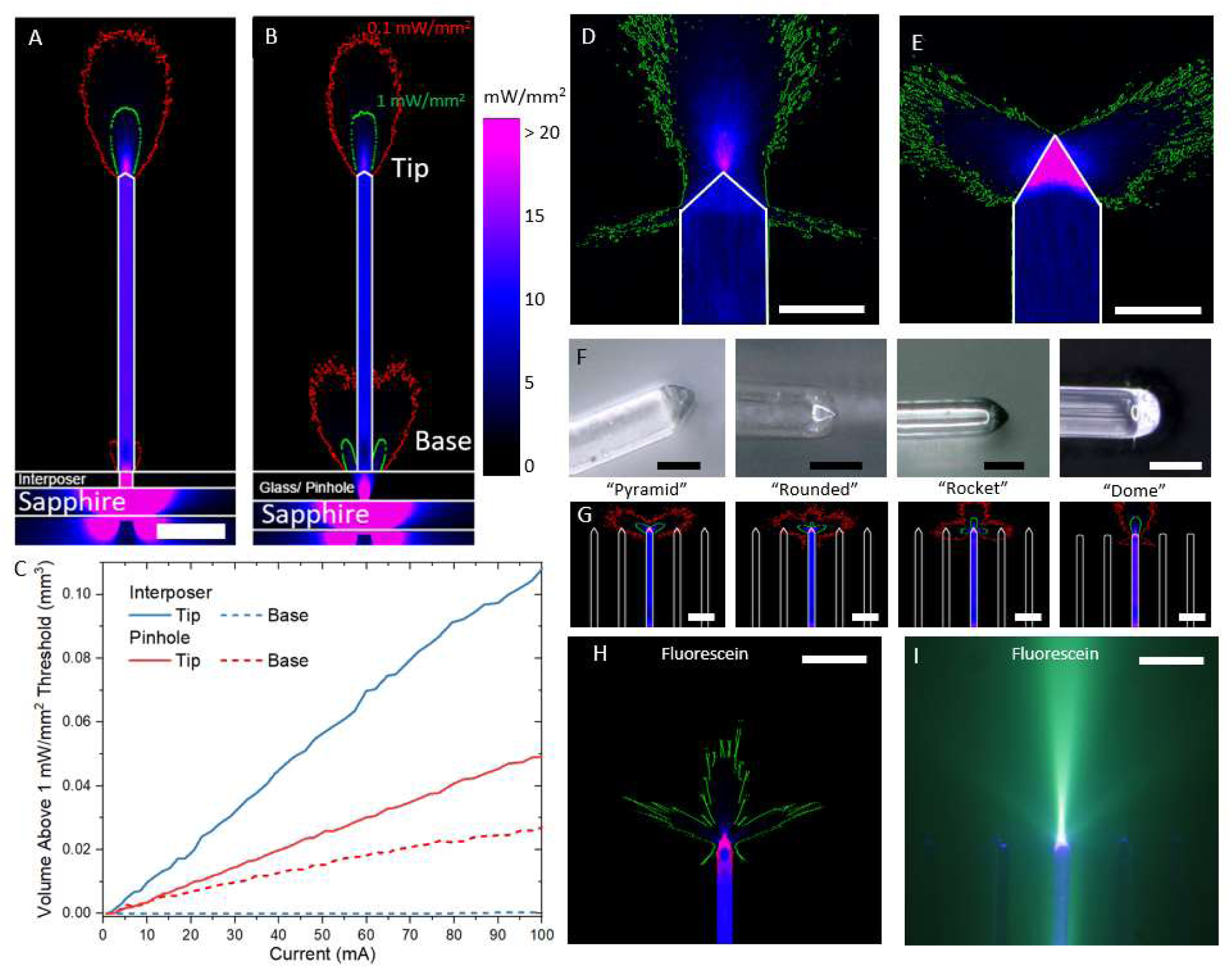
**A** Cross-section of the modelled light output in cortical tissue from the UOA (30° tip angle) with the new optical interposer. The green contour is the 1 mW/mm^2^ level, and the red contour is 0.1 mW/mm^2^. The scale bar is 400 µm. **B** The modelled light output in brain tissue from the UOA with a pinhole layer to limit stray light [27]. **C** The volume of excitation for a given current for both the optical interposer UOA and pinhole UOA. **D-G** Alternative tip shapes are possible allowing various light profiles to be coupled into the brain. Low-angled tip (30°) gives the highest peak irradiance (see **A** and **B** with a close-up in **D**), while a 60° angled tip (panel **E**) distributes light laterally. This is the tip shape that was used in vivo. **F** Images of the different tip shapes can be realised by extending the annealing time of the needles during array fabrication. Scale bars in **D-F** are 100 µm**. G** Optical modelling of light emission and scattering in brain tissue, for each of the different tip shapes in **F**, scale bar is 400 µm. Optical modelling (**H**) and device image (**I**) of the light emission in a fluorescein solution, for comparison. The scale bar is 400 µm.

Finally, the die can be singulated with a dicing saw into individual UOAs, followed by a final cleaning step to remove any contamination from mounting the die during dicing. A completed needle array is shown in Figure 1E. Final needle lengths were 1.5 mm. Needles of between 0.5 and 2 mm can be achieved by modifying the dicing and annealing steps[23].

### 2.3 Fabrication – µLED array

The µLED array is fabricated on a commercial InGaN/GaN wafer. The III-nitride materials are grown on a c-plane (0001) 2-inch sapphire substrate. The GaN layers consist of an undoped GaN buffer layer, an n-type GaN layer and a multi-quantum well layer consisting of layers of InGaN and GaN designed to emit light at 450 nm. The MQW region is covered by an electron-blocking layer (*p*-type AlGaN), to help confine electrons to the MQW region.

The fabrication process flow of the µLED is detailed in [27] and discussed briefly here. Initially, electron beam evaporation is used to deposit a 100 nm-thick palladium current spreading layer on top of the *p*-type region. To ensure a good ohmic contact between the palladium and GaN the device is annealed at 400 °C for 3 minutes in a N_2_ ambient. An inductively coupled plasma (ICP) process (Ar:Cl, 10:30 sccm flow) is used to etch mesa structures in the *p*-type layer (masked by a 300 nm PECVD silicon oxide layer) exposing the *n*-type GaN layer and defining the μLED pixels. A Ti:Au metal stack (100:300 nm) is sputter deposited to create tracks connecting to the *n*-type region. A thin film passivation layer (1000 nm PECVD silicon dioxide) covers these tracks and *n*-type regions, with vias etched (Ar-fluoroform mixture in a reactive ion etch tool) to open electrical contact sites to the p-mesa. A second metal layer is then deposited to connect to the *p*-type regions (Ti:Au 50:300 nm). A further 300 nm of PECVD silicon dioxide protects the surface, with a further RIE step to open the bond pad sites around the periphery of the device (see Figure 1C). Bond pads have an additional Ti:Pt:Au (100:200:400 nm) layer sputter deposited to improve the wire bonding yield.

### 2.4 Device Fabrication – Integration and Encapsulation

Integration of the device was completed using a flip-chip bonder (Fineplacer pico-2, Fintech). The glass needle array was held in a custom-made holder and the µLED array was brought to within 10 µm of the UOA. Imaging was used to ensure accurate alignment. An underfill capillary gap-filling method was used to dispense the UV curable glue (Norland 61) as this ensured no air bubbles and gave the best overall alignment. Optical modelling was used to determine the misalignment tolerance (Appendix 1). This indicates that there is little loss in optical coupling efficiency for misalignments up to 20 µm.

Developing an electrical connection scheme to address each µLED separately is challenging in terms of track routing and number of bond pad sites. Therefore, a matrix-addressing scheme was adopted. In this approach, all pixels along one column share a common anode (*p*-contact) and all pixels along one row share a common cathode (*n*-contact). Therefore, 38 connections are required, 19 anodes and 19 cathodes, which simplifies the electronic driver scheme dramatically compared to individually addressed µLEDs. This approach allows commercial LED current drivers to be employed, and reduces the number of connections, at the cost of limiting the available patterns that can be displayed. For example, individual µLEDs, horizontal or vertical lines and rectangles/squares are possible. Diagonal light patterns and simultaneously displayed horizontal and vertical lines are not possible unless pulse width modulation schemes are used to realise these restricted patterns (see supplementary movie 1).

Each of the 38 connections is linked to LED driver circuitry through insulated gold wire bonds (25 µm diameter and up to 10 cm long). The bond pads and wire bonds were then coated in silicone and Parylene-C was deposited to further encapsulate the device. The finalised device, with a single needle site switched on, is shown in Figure 1A.

### 2.5 Electrical and Thermal Characterisation

The output from each µLED was characterized electrically and optically, before and after final integration, using a programable power supply (B2901A, Keysight Technologies) and optical power meter (S120VC, Thorlabs). The optical power meter has a limited collection aperture; therefore, a geometric correction factor was used to calculate the total optical power emitted by the µLED. The correction was benchmarked against an integrating sphere measurement. The thermal performance of the device was measured in air using an infrared camera (SC7000, FLIR). Thermal modelling using finite element analysis (FEA) software (COMSOL Multiphysics) allowed the thermal measurements in air to be related to device performance *in vivo*. The material property inputs to the model are included in Table 1. It was assumed that 90% of the electrical power was converted to heat in the µLED structure. A brain perfusion rate of 0.5 L/kg/min was included in the COMSOL model [33]. µLED drive currents (0.5-100 mA), pulse widths (1-100 ms) and frequencies (1-100 Hz) were modelled, as were multiple (simultaneously illuminated) µLEDs. To allow the accuracy of the model to be assessed, the temperature at the top of the device encapsulation was measured using a thermal camera during *in vivo* experiments.

**Table 1:**
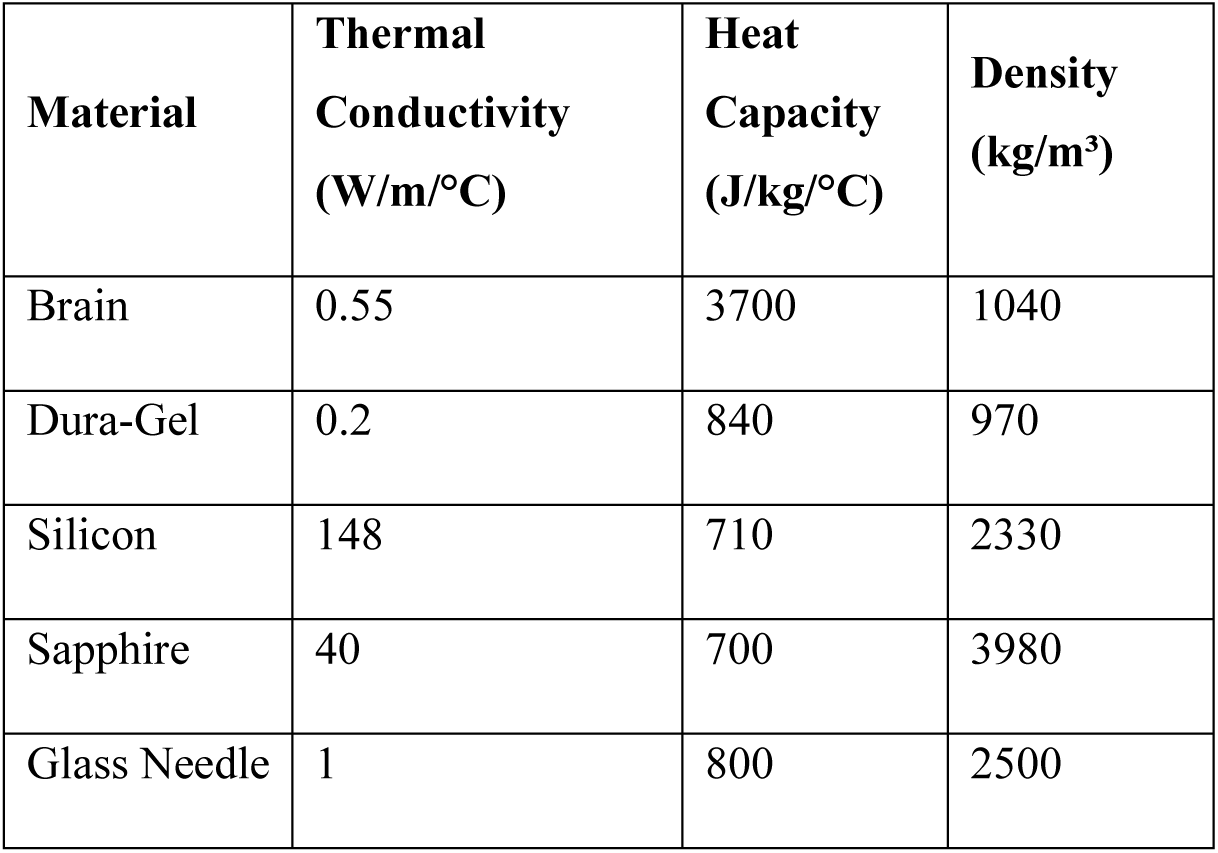
Thermal properties used in the FEA simulations.

### 2.6 In vivo testing

The device was tested in the left hemisphere of one sufentanil-anesthetized adult female cynomolgus monkey (*Macaca Fascicularis*). To restrict expression of the excitatory opsin channelrhodopsin-2 (ChR2) and the red reporter protein tdTomato to excitatory neurons, the primary visual cortex (V1) was injected with a mixture of Cre-expressing and Cre-dependent adeno-associated viral vectors carrying the genes for ChR2 and tdTomato (AAV9.CamKII.4.Cre.SV40 and AAV9.CAG.Flex.ChR2.tdTomato, Addgene). Following a post-injection survival period of 9 weeks, to allow for opsin expression, the animal was anesthetized and prepared for an acute terminal experiment. A full description of the procedure is available elsewhere [30]. The UOA was inserted into the opsin/tdTomato-expressing region of V1 to a depth of approximately 1 mm (the curvature of the visual cortex causes different needle tips to be at different depths) using a high-speed pneumatic electrode inserter system (Blackrock Neurotech). Electrophysiological recordings were made with a 24-channel linear electrode array (LEA), with an electrode spacing of 100 µm and a 300 µm distance from the tip to the first electrode (V-Probe, Plexon). The LEA was inserted into the cortex, close to the UOA, in two separate penetrations. The first penetration, P1, was approximately 1 mm from the UOA (nearest µLED - row 8, column 1), inserted to a depth of 2.4 mm. The second penetration, P2, was inserted to a depth of 2.6 mm and angled towards the UOA so that the deeper contacts were approximately 800 µm from the UOA (nearest µLED - row 5, column 1), while the more superficial contacts were approximately 900 µm from the UOA (nearest µLED-row 6, column 1). A 128-channel recording system (Cerebus, Blackrock Microsystems) was used to record the electrical data. Data were sampled from the 24 channels at 30 kHz. A silicone gel (Dura-Gel, Cambridge NeuroTech) was applied across the exposed cortical surface[34], underfilling the UOA. Finally, GELFOAM (Pfizer) was placed over the implant site to protect the exposed scalp tissue, and periodically soaked with saline. Each experiment had the same optical stimulation protocol: Light was pulsed at 5 Hz, with 100 ms pulses for a duration for 1 s, followed by a 1.5-21 s inter-trial interval (longer inter-trial intervals were used at the highest photo-stimulation intensities). For each experiment, approximately 5 minutes of data was recorded giving 30-45 trials, each with five 100ms µLED pulses. The µLED drive current, location and number of illuminated µLEDs was changed between experiments. All procedures conformed to the US National Institutes of Health Guide for the Care and Use of Laboratory Animals and were approved by the University of Utah Institutional Animal Care and Use Committee.

### 2.7 In vivo data analysis

Multi-unit activity (MUA) is a high-frequency signal band, often used as a measure of the spiking activity of many neurons in the vicinity of a recording electrode and was clearly observed in response to UOA activation of ChR2-expressing neurons. To assess the level of activation the data was first processed by employing a band stop filter (2^nd^ order Butterworth) to remove 60 Hz interference from mains power. The multi-unit activity was quantified by further filtering using a band pass filter from 0.3 kHz to 6 kHz and the absolute value of this signal on each electrode was taken as a function of time. The data from multiple trials was then aligned based on the µLED pulse time and averaged. Low-frequency activity or Local Field Potential (LFP) data can also be extracted using a band pass filter between 1 and 100 Hz. The Current Source Density (CSD) was calculated from the LFP data using a kernel CSD method (kCSD_Matlab [35]). The CSD reveals the location of current sinks (neuron depolarisation) and sources (return currents) throughout the cortical depth. To average the signal from multiple trials, the data was aligned based on the turn-on time of the µLED pulse.

## 3. Results

### 3.1 Optical Performance

The modelled optical output of the UOA device, with optical interposer, is shown in Figure 2A. The device has a modelled optical efficiency of 0.4% (the previously reported pinhole device had a modelled efficiency of 0.22%[27]). Due to the silicon optical interposer the stray light emitted at the base of the needles has also been reduced by a factor of 12 compared to the previous device (Figures 2 A and B). Figure 2C shows the modelled volume of tissue illuminated with an irradiance of greater than 1 mW/mm^2^ for both the new interposer UOA and the former pinhole device. The interposer device gives an expected volume above a 1mW/mm^2^ threshold of 2.1 times that of the pinhole device.

The optical models were further used to determine the optimal tip angle for the needle device, with a 30° tip outcoupling most light. However, this is not ideal for device implantation, needles with larger tip angles minimise the insertion force required to penetrate the brain and reduce trauma and vascular damage [36]. Optical modelling indicates that low tip angles (30°) give deep illumination, while large tip angles (>60°) give a lateral profile more suited to laminar optogenetic excitation (Figures 2D and E). Sideways emission is caused by reflected rays, from the glass-brain interface, being out-coupled at the opposite facet. Above a tip angle of 60°, all the light will be directed laterally due to total internal reflection. The device used for the *in vivo* study had a tip angle of 60°.

The tip geometry can be controlled by the bevel angle of dicing blades, and the heat treatment used to smooth the optrode surface [19]. The annealing step causes a rounding of the corners at the tip of the device. If the annealing time is increased the tip shape becomes more rounded, starting with rounding of the sharp corners of the tips, before transitioning into a rocket tip shape and then into a hemispherical dome shape (Figure 2F). This had a similar effect to altering the tip angle and can give the device user a pre-determined choice of optical profile depending on the brain region they are targeting (Figure 2G).

Figures 2H and I show the modelled and imaged optical output from a test device in a fluorescein solution. This confirms that the optical model captures the light emission profile from the device.

Multiple, simultaneous illumination sites were also modelled (Appendix 2), with the magnitude of each µLED output given by measured values for the fabricated device. In this device architecture iteration, the interstitial sites were not coupled to TSVs, as we adopted an iterative approach to device design and had concerns on over-complicating the first fabrication run. These concerns proved unfounded, and we have since made devices with operational interstitial sites (Appendix 2), though these have not been tested *in vivo*. Modelling of the interstitial sites indicate that a distance of up to 1 mm, from the surface of the cortex, can be illuminated at an irradiance above 1 mW/mm^2^.

Figures 3A and B show the average irradiance output of the device when each µLED is driven at 20 mA (60° tip shape). Since a tip angle of 60° was used here, there is a lateral outcoupling of light and no near-field focus (as previously reported [27] and shown in Figure 2D). For this reason, we do not quote the peak irradiance, but instead give the average irradiance across the emitting area of the tip, as shown in Figure 2E. The non-uniformity in figure 3A and B in average irradiance is a result of fabrication variances across each µLED. The array uniformity can be improved by adjusting the current/voltage on a pixel-by-pixel basis. However, the maximum tip irradiance will be set by the least bright µLED included in the adjustment (3 mW/mm^2^ in Figures 3C and D). In this case, 94 out of 100 µLEDs will emit between 3 and 3.4 mW/mm^2^. This could be improved further by replacing the voltage source used here with a programmable current source with µA steps between current levels. The current and voltage required to achieve uniform illumination are shown in Appendix 3.

**Figure 3:**
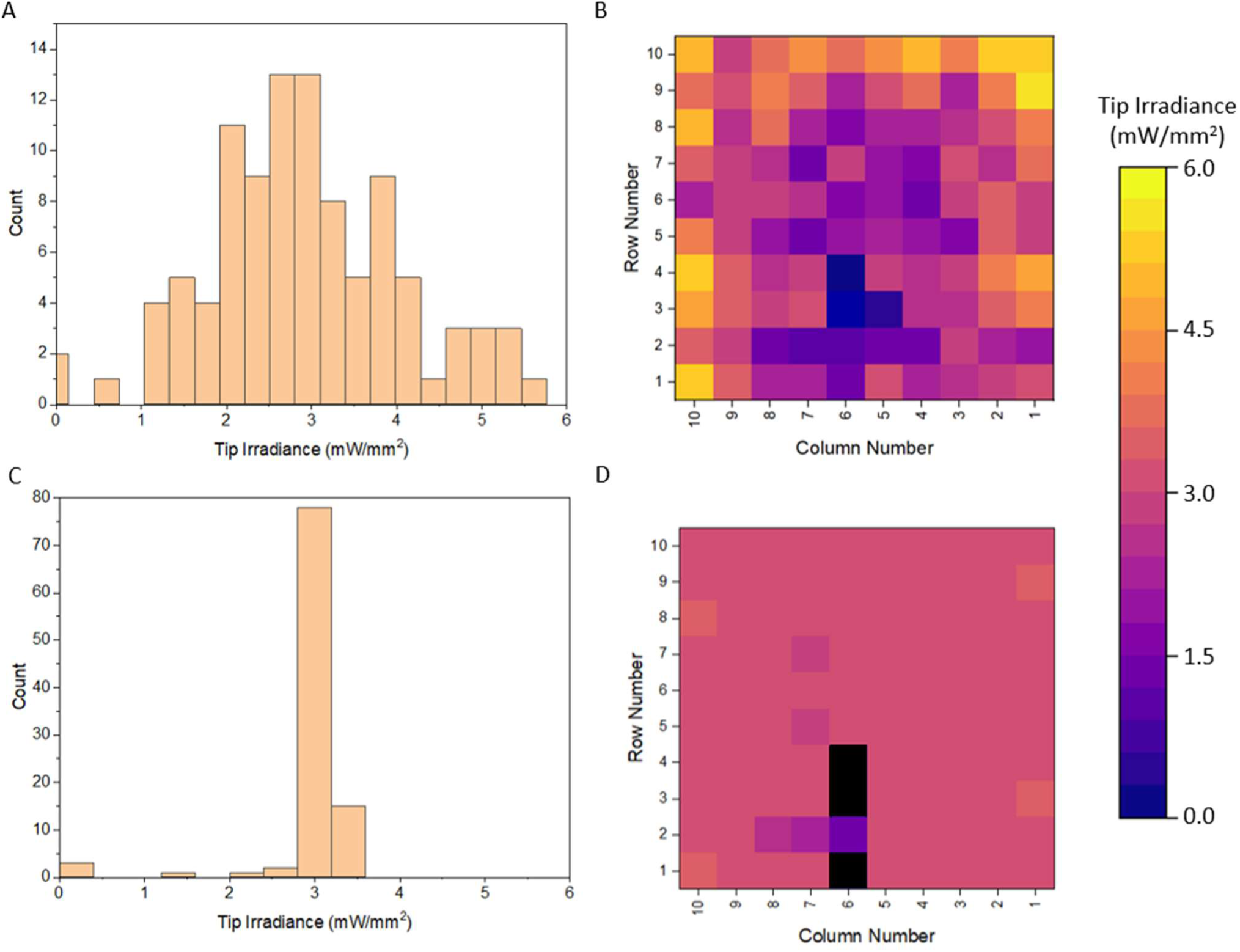
**A** Histogram of the irradiance at the tip of the needles when the µLEDs were driven at 20 mA. **B** Map of the irradiance at the tip of the needles at 20 mA drive current. **C** Histogram of the irradiance at the tip of the needles when each µLED was driven with a programable current source, aiming for an irradiance of 3 mW/mm^2^. **D** Map of the irradiance at the tip of the needles when the device was operated with a programable current source, aiming for irradiance of 3 mW/mm^2^. Black pixels did not have a stable optical output and were excluded from the map.

### 3.2 Thermal Performance

It has been observed that small temperature deviations of ∼1°C can change the behaviour of neurons [28]. Temperature changes of this magnitude can also drive changes in behaviour [29]. Previously the thermal constraints when the device was fully inserted were explored and the temperature at the tips of the needles where light is emitted was reported [27]. However, thermal modelling indicates that the temperature at the base of the needles will get significantly warmer and could potentially alter the behaviour of superficial neurons. Here the operating range for the µLEDs is extended by partially inserting the array and underfilling it with a thin layer (∼500 µm) of silicone (Dura-Gel) between the brain and the device backplane. The primary purpose of this layer is to prevent the surface of the brain from drying out during experiments [34]. However, this thermally insulating layer significantly reduces temperature increases at the cortical surface compared to direct contact between the device and cortical tissue. Figure 4A shows the device implanted into the monkey area V1. The Dura-Gel is transparent and not visible in the image. The GELFOAM (labelled) is placed on top of the Dura-Gel and used to prevent the tissue around the scalp from drying out. This is periodically soaked in saline throughout the experiment. Figure 4B shows the maximum temperature recorded using a thermal camera while the device was operated with 10 simultaneously activated µLEDs, each operated at 5V (∼30 mA), 100 ms pulses at 5 Hz for 1s followed by 10 s off. This measurement gives a 4.1°C temperature rise at the top of the Dura-Gel, which coated the device. From thermal modelling, this surface temperature corresponds to a brain surface temperature increase of 1°C. Figure 4 C shows a schematic of the geometry of the thermal model, the highlighted points are the points where the modelled temperature is recorded: Green – Air/Dura-Gel boundary; Red – Interposer/Dura-Gel boundary; Black – Cortex/Dura-Gel boundary; Blue – tip of the needle. Figure 4D shows the temperature of various points on the device for the specific stimulation pattern that was used in the *in vivo* experiments (100 ms pulses at 5 Hz for 1 s followed by 10 s off.). The colours of the lines correspond to the points highlighted in Figure 4C. The green line is at the Dura-Gel/Air boundary and corresponds to the measurement point using the IR camera (Figure 4B), which reaches a maximum temperature approximately 2 s after the start of the stimulation trial. The difference between these two values (modelled: 3.1 °C increase, measured: 4.1 °C increase) is likely due to the estimation of the Dura-Gel thickness (taken to be ∼300 µm) that lies above the device. Further stimulation protocols are compared in Appendix 4, showing good agreement in each case. Although the device itself can increase in temperature by several degrees during operation, the modelling indicates that the surface of the cortex (black line in Figure 4D) remains just below 1 °C with the tip of the needle (blue line) showing no temperature change for this specific stimulation pattern.

**Figure 4:**
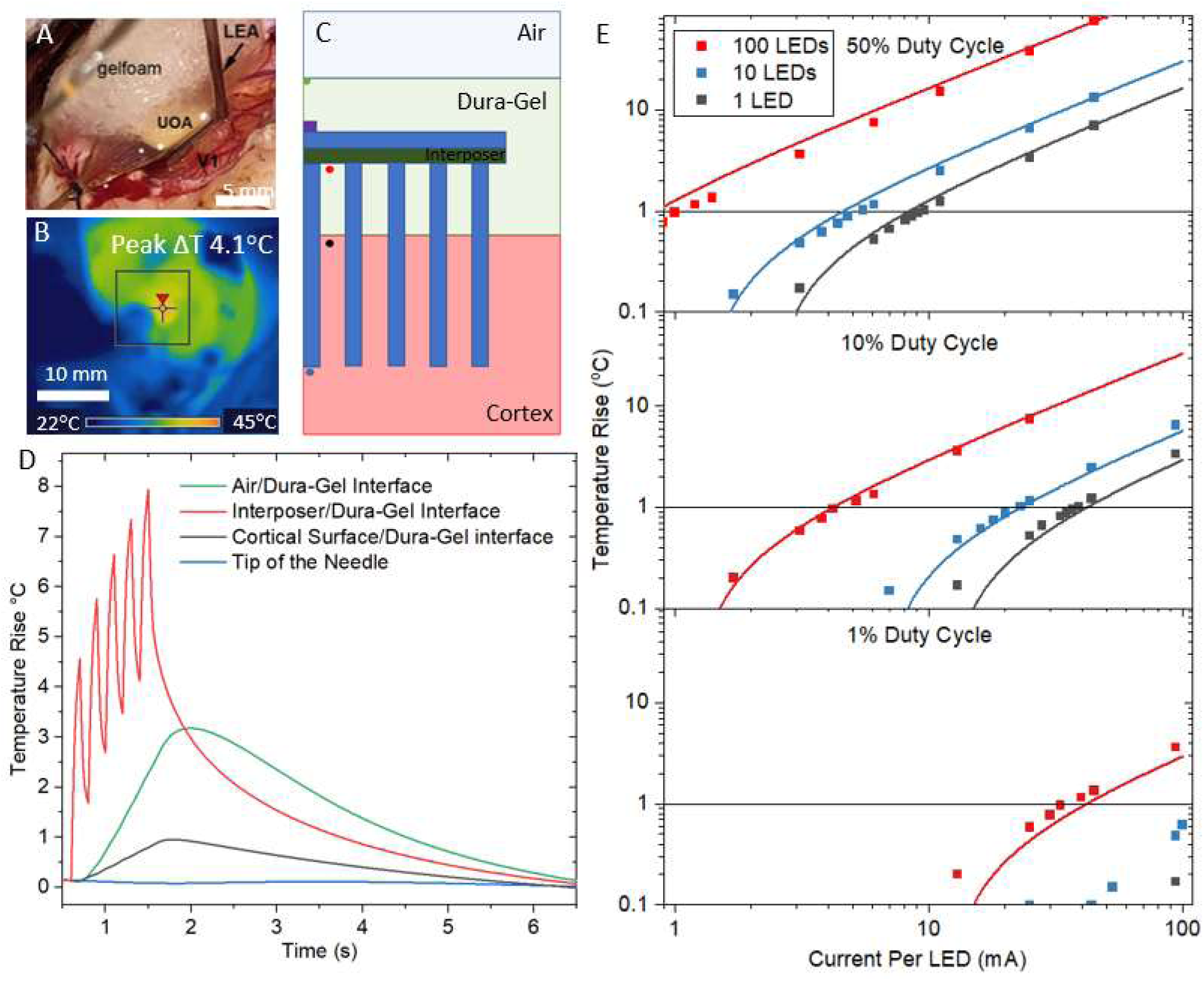
**A** Image of the in vivo experimental setup. The UOA and LEA are labelled. After insertion, the device and brain surface are coated with Dura-Gel to prevent the surface of the cortex from drying out (not labelled). GELFOAM (labelled) is applied on top of the device to further prevent tissue drying. **B** Thermal camera image of the device in operation (10 simultaneously operated µLEDs at 5V (∼30 mA), 100 ms pulse at 5 Hz for 1 s followed by 10 s off). The peak measured temperature rise was 4.1°C above the baseline measurement. **C** Schematic showing the simplified structure that has been modelled. The coloured dots correspond to the colours of the lines on the graph in **D**. **D** Thermal modelling data generated using the specific excitation protocol of 10 simultaneously illuminated µLEDs (5 V at ∼30 mA), 100 ms pulse at 5 Hz for 1 s followed by 10 s off. The predicted temperature rise at the air/Dura-Gel interface compares well with the thermal camera measurement. **E** The modelled steady state temperature systematically analysed as a function of µLED currents, duty cycles and the number and pattern of µLED illuminated. The plotted line is taken from Equation 1.

Figure 4E shows a more general case of continuously pulsing the µLEDs at various duty cycles, currents, and number of activated µLEDs. The quoted temperature here is for the point at which the device reaches a steady-state equilibrium temperature (i.e., where thermal generation by the □LEDs is balanced by heat dissipation in tissue and through the device). This occurs after approximately 35 s of continuous operation for any given duty cycle. From these results, a linear fit to the data gives a formula which can provide the approximate temperature increase in the brain (ΔT, °C) for a given duty cycle (D), number of µLEDs (N_LED_), drive current (I_LED_, mA) and duration of the pulse train (t, s) (Equation 1).

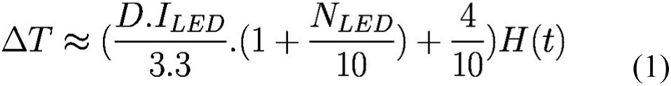

H(t) is a Hill growth function given by H(t) = t^1.6^/(6+t^1.6^). For example, setting the maximum temperature rise to 1°C and calculating for 10 µLEDs, operated at a 10% duty cycle, give a µLED current limit of 17 mA. This current is sufficient to give an irradiance of 2.5 mW/mm^2^ across the emitting surface of the 60° tipped needle.

### 3.3 In vivo Testing

ChR2 and tdTomato were expressed in the macaque visual cortex (V1 and V2) via a mixture of cre-expressing and cre-dependent adeno-associated viral vectors. The UOA was inserted into V1 (Figure 5A), an area of high expression. that was checked post-experiment through histological imaging of tangential sections (Figure 5 B). After a period of recovery, the LEA was inserted and electrical activity recorded while the UOA scanned through stimulation protocols.

**Figure 5:**
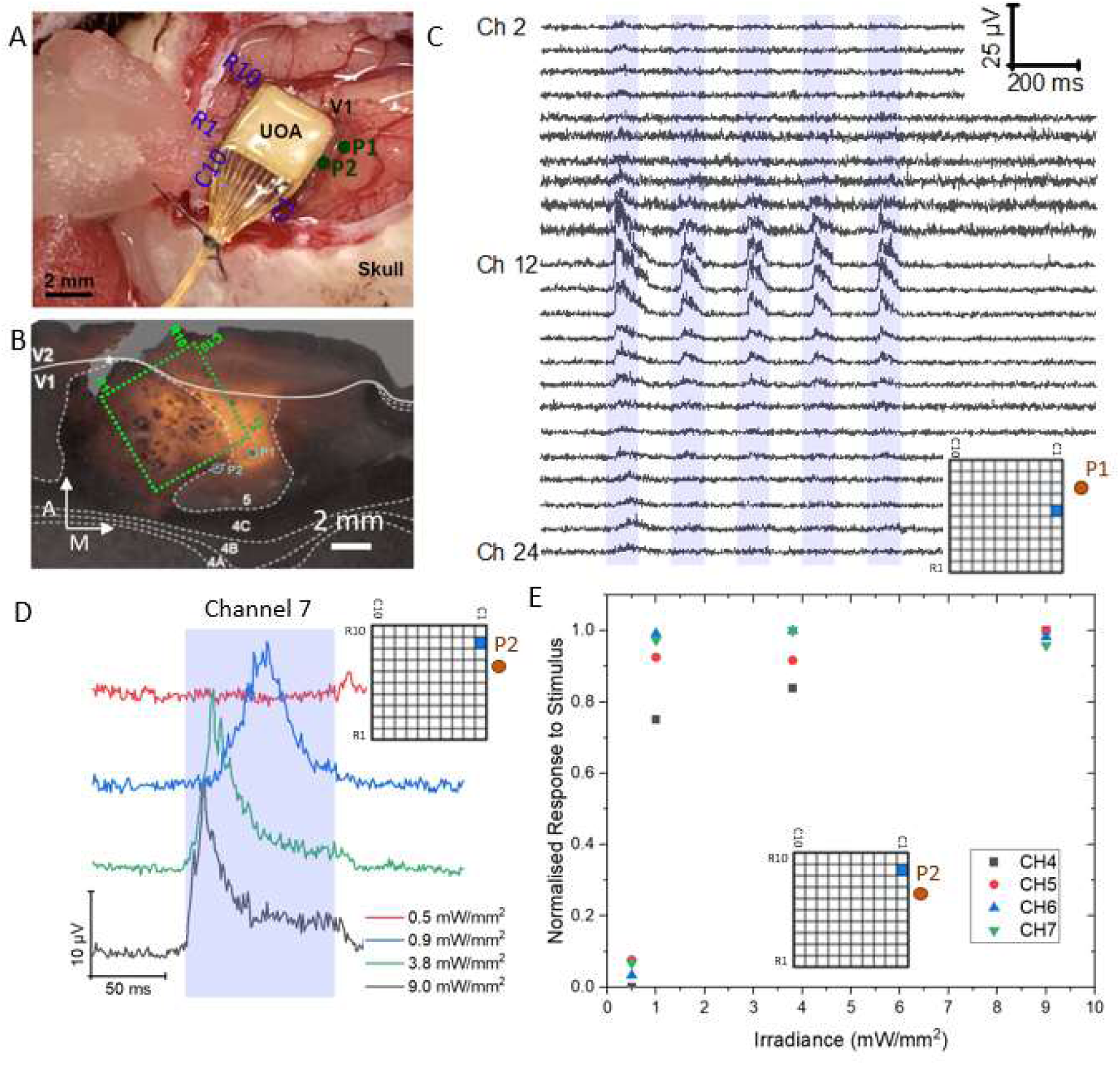
**A** Image of a UOA implanted in the V1region of the primate visual cortex. µLED row and column numbers are labelled. The LEA was inserted at two locations P1 and P2. **B** An epifluorescence microscope image of a Cytochrome-oxidase stained and tdTomato fluorescent protein tangential section through V1 and V2. The section is approximately 900 µm deep. The solid white line marks the boundary of V1 and V2. The dashed white contours delineate layers in V1. The location of the UOA is shown with a dashed green line and the LEA positions are also marked. The orange emission images the location of the tdTomato co-expressed with ChR2. A: anterior; M: medial **C** MUA recordings from a single trial with the LEA positioned at P1. µLED Column 1 Row 6 was illuminated with a drive current of 24 mA, giving a peak irradiance of 3.1 mW/mm^2^. The blue shading indicates when the µLED was on. **D** Average MUA recordings from LEA channel 7 at P2 while µLED Column 1 Row 9 was driven at currents of 0.6 mA, 1.4 mA, 11.4 mA and 44.1 mA (0.5 mW/mm^2^, 0.9 mW/mm^2^, 3.8 mW/mm^2^ and 9 mW/mm^2^). Each trace represents the average of 150 trials. **E** Dose-response curve plotted as the normalized response as a function of irradiance for µLED Column 1 Row 9 and LEA at P2. The MUA was taken from the 4 electrodes showing the greatest increase in activity.

For each LEA insertion, a strong µLED dependant increase in MUA could be seen at the approximate location of the tips of the UOA (Figure 5C), channel 12 in P1 and channel 7 in P2, this corresponds to a cortical depth of 1.1 mm, aligning with layer 4C in the visual cortex. The average response of 150 µLED pulses on electrode channel 7, while µLED C1 R9 was illuminated at different irradiances, is shown in Figure 5 D. The irradiance is calculated as the average irradiance across the area of the tip where light is emitted. This allows us to create a dose-response curve (Figure 5E). The onset of optogenetic excitation for this limited experiment was an average tip irradiance of 0.9 mW/mm^2^.

In Figure 6, the LFP and current source density (CSD) of the optogenetic response (as recorded by the LEA at insertion point P1) is shown and analysed as a function of which µLED was activated. The motivation was to assess the distinctiveness of the neural response for different µLED positions. In Figure 6A the average LFP response from 31 trials when µLED C1 R4 was switched on with an average tip irradiance of 11 mW/mm^2^ is shown, this shows a strong positive deflection around channel 12 approximately 50 ms after the onset of the µLED illumination. In Figure 6B the CSD calculated from the data in Figure 6A is shown. This shows a strong current sink in the layer where the tips of the needles were located (point of highest irradiance). The minimum value of the corresponding sink is approximately 75 ms after the onset of the 100 ms duration µLED illumination.

**Figure 6:**
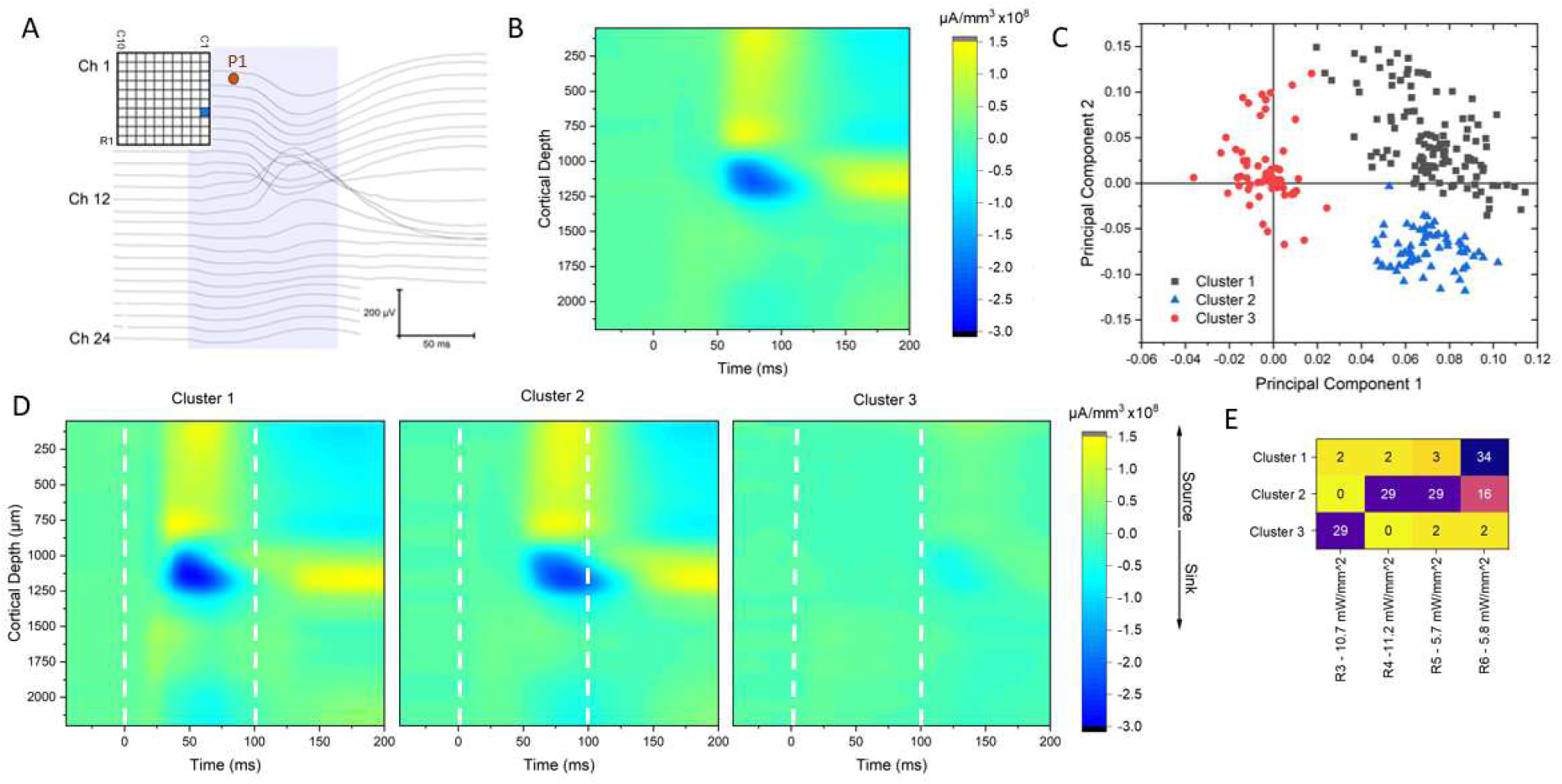
**A** LFP at P1 while µLED Column 1 Row 4 was operated at 100 mA, 11.2 mW/mm^2^ (average of 31 trials). The blue shading indicates when the µLED was on. Activation at the depth of the UOA tips is clearly visible. **B** CSD plot while µLED Column 1 Row 4 was operated at 100 mA, 11.2 mW/mm^2^ (average of 31 trials), calculated from the LFP example in A. The µLED switched on at time 0 for 100 ms. **C** PCA of the CSD from electrical recordings at P1 from 148 individual trials using µLEDs Column 1 Rows 3-6 operated at 100 mA, Irradiance of between 5.7 and 11.2 mW/mm^2^. The PCA was clustered based on the first 5 principal components (which contain >80% of the data variability), forming 3 distinct clusters. Shown here is a scatter plot of the first two principal components. **D** CSD Plots for each of the 3 clusters in C, showing that each cluster creates its unique pattern of electrical activity. **E** Correlation matrix indicating that cluster 1 corresponds to µLED R6, Cluster 2 contains µLEDs R4 and R5 and Cluster 3 contains µLED R3, demonstrating that μLED-to-LEA proximity dictates the recorded neural pattern.

To demonstrate how different μLED sites can induce different patterns of LFP response, the individual trials from 4 different µLEDs were taken (C1-R3, C1-R4, C1-R5 and C1-R6), the CSD for each trial was calculated and a principal component analysis (PCA) was used to determine the statistical variance between the CSD plots, by projecting the data into principal component space. Similar LFP/CSD responses should cluster into distinct regions of the phase space and the correlation of this clustering with μLED position will give an indication as to the distinctiveness of the induced neural activity. K-means clustering was used on the first 5 principal components to define the clusters (containing ∼80% of the data variability). The data clusters into 3 distinct populations (figure 6C), each showing differing neural responses. When plotting the average CSD for each cluster (figure 6D): cluster 1, shows the largest response with the shortest latency to the deepest current sink (50 ms); Cluster 2, has a latency that increases to 90 ms and a reduction in the current sink depth; cluster 3 shows a greatly reduced response and an increased latency (125 ms) to the minimum current sink. Which µLED is responsible for which cluster is examined in the correlogram in Figure 6E. Cluster 1 has 41 entries, 34 of which are from µLED C1 R6 stimulation (the nearest to the LEA). Cluster 2 has 74 entries, mostly attributed to µLED C1 R4 and R5 (29 entries each). Cluster 3 has 33 entries, with 29 of them from µLED C1 R3 (the µLED furthest from the LEA). This analysis demonstrates that there is a statistically significant difference in the optically induced cortical activity pattern from each µLED site. As the UOA stimulation sites becomes more distant, the latency of the current sinks, detected on the LEA, increases. In Appendix 5 the statistical difference between each of the clusters that were identified in the PCA analysis is quantified.

## 4. Discussion

The UOA with integrated silicon optical interposer presented in this work had an optical efficiency of 0.4%. Although this appears low, it is 80% greater than the previously reported efficiency from the UOA with a pinhole layer [27]. This increase in efficiency extends the range over which the device can operate without exceeding thermal limitations. This means an increased irradiance and so an increased volume above a given threshold (e.g., 1 mW/mm^2^ for ChR2). It could also mean an increased stimulation duty cycle to better match natural neuronal activity. Implantable µLED probes can achieve >10% optical efficiency; however, the µLED source is in contact with delicate and temperature sensitive neural tissue, meaning that thermal constraints also limit the maximum optical output [14, 15]. Further increases in the optical efficiency of the UOA could be achieved by filling the interposer holes with a material that matches the refractive index of the glass needles, further thinning the interposer layer, designing a top emission µLED device or the integration of micro-lenses to improve optical coupling [37].

Stray light, and so optical crosstalk between sites, has also been reduced by more than an order of magnitude through the introduction of a silicon optical interposer. In the previous version of the UOA, a pinhole layer was used to reduce stray light. While this was effective when compared to the case without a pinhole layer, there was still a volume of tissue (∼0.007 mm^3^ when the µLED is operated at 20 mA) at the surface of the cortex that would be above 1 mW/mm^2^ irradiance. This caused some ambiguity about the region where an optogenetic signal was being generated. The UOA with interposer achieves illumination at the tip of the needle with no stray light (above the stimulation threshold) in the superficial layers of the cortex. This new design removes uncertainty in the stimulation region and allows for a greater scope of *in vivo* experiments. Further improvements to remove any residual stray light can also be considered, including coating the base of the needles with an optical absorber.

The output and beam profile from the needle tip can also be modified by changing the bevel angle of the dicing blade or annealing time during the fabrication of the glass needles (figure 2) [23]. If deep illumination is required, a shallow tip angle (∼30°) should be used, while if a laminar illumination profile is preferred, a tip angle of ∼60° is required.

Both the depth and lateral resolution of optogenetic stimulation were investigated as part of the *in vivo* studies. Figures 5C and 6A both show clearly defined neuronal activity, correlated to the optogenetic stimulation in the location of the UOA needle tips. The depth distribution of evoked activity spans across ∼6 electrodes, 600 µm. No activity (correlated to the optogenetic stimulation) was observed in the most superficial recording sites (less than a depth of 500 µm) in either case. The extent of the optogenetically driven activity corresponds well with the optical modelling in figure 2F and supplementary figure A2. Lateral resolution is more difficult to quantify with the current experimental methodology. However, the analysis in Figure 6 indicates that each needle site is producing a different pattern of neuronal activity, suggesting that higher resolution devices may add functionality.

Increasing the transverse resolution, by reducing the pitch of the needles, is challenging with the current fabrication approach, particularly since the needle diameter would also likely have to be reduced to minimise brain volume displacement. Thinner needles will reduce optical coupling efficiency and are more likely to break during fabrication, affecting device yield and requiring process optimisation. Alternate methods for producing transparent needle arrays are being investigated including two-photon polymerization [38].

Thermal modelling of the device indicates that there is a broad range of drive currents and duty cycles which will not increase the temperature above a 1°C limit. In the previous work [27], the UOA was considered as an isolated device inserted into tissue. In this work, a full system approach is used. A silicone gel (Dura-Gel, Cambridge Neurotech) layer, whose primary role is to prevent the delicate cortical tissue from drying out [34] is found to also act as a thermal barrier and enhances the operational range of the device (when a single μLED is operated at 20 mA with a 10% duty cycle the temperature increase at the cortical surface has been reduced from 2 °C to 0.9 °C). The bond wires used to drive the µLEDs were found to act as a route for heat conduction away from the device, thereby reducing the thermal energy reaching the cortical surface. This points to further improvements that could be made to the device, including an improved thermal barrier between the device and the cortical surface, and intentionally using integrated approaches, that optimize thermal conduction away from the device.

## Acknowledgements

We thank Julian Haberland and Christine Kallmayer at the Fraunhofer-Institut für Zuverlässigkeit und Mikrointegration for µLED-to-interposer bonding. This work was supported by the NIH BRAIN Initiative through Grant No. U01 NS099702. KM was supported by the Royal Academy of Engineering under the Chair in Emerging Technologies scheme.

## Appendix 1

Misalignment between the µLED array and UOA can be optically modelled to determine its effect on light coupled into the needle and stray light (figure A1). From this analysis, the misalignment tolerance between the µLED array and interposer needle array is approximately 20 µm.

**Figure A1:**
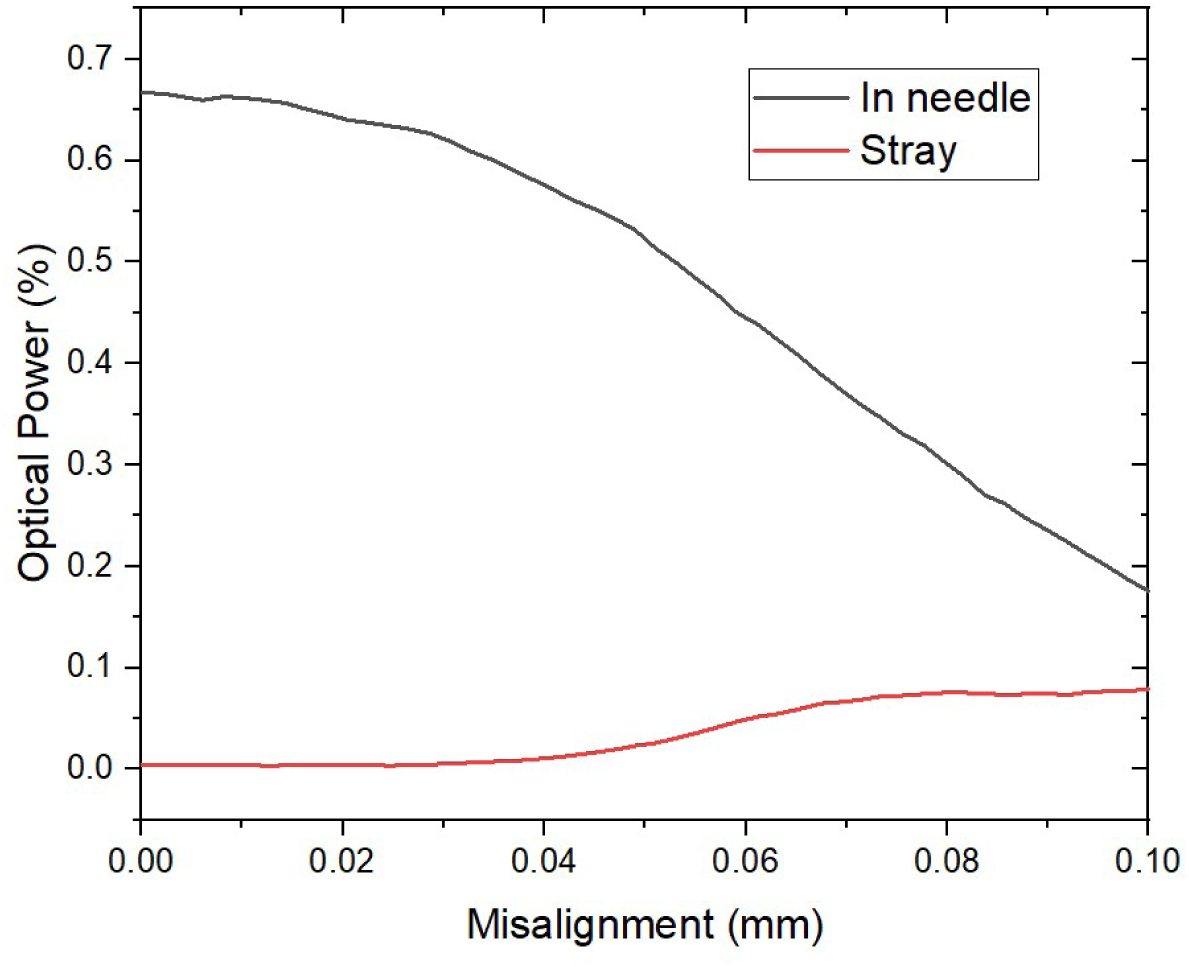
Misalignment between the µLED array and needle array versus the optical power coupled into the needle and stray light.

## Appendix 2

The emission from multiple simultaneously illuminated sites can be modelled by taking the emission from a single site, duplicating it, shifting it in x or y and summing it with the original model of a single emission site. Figure A2 shows the emission from several sites illuminated simultaneously.

**Figure A2:**
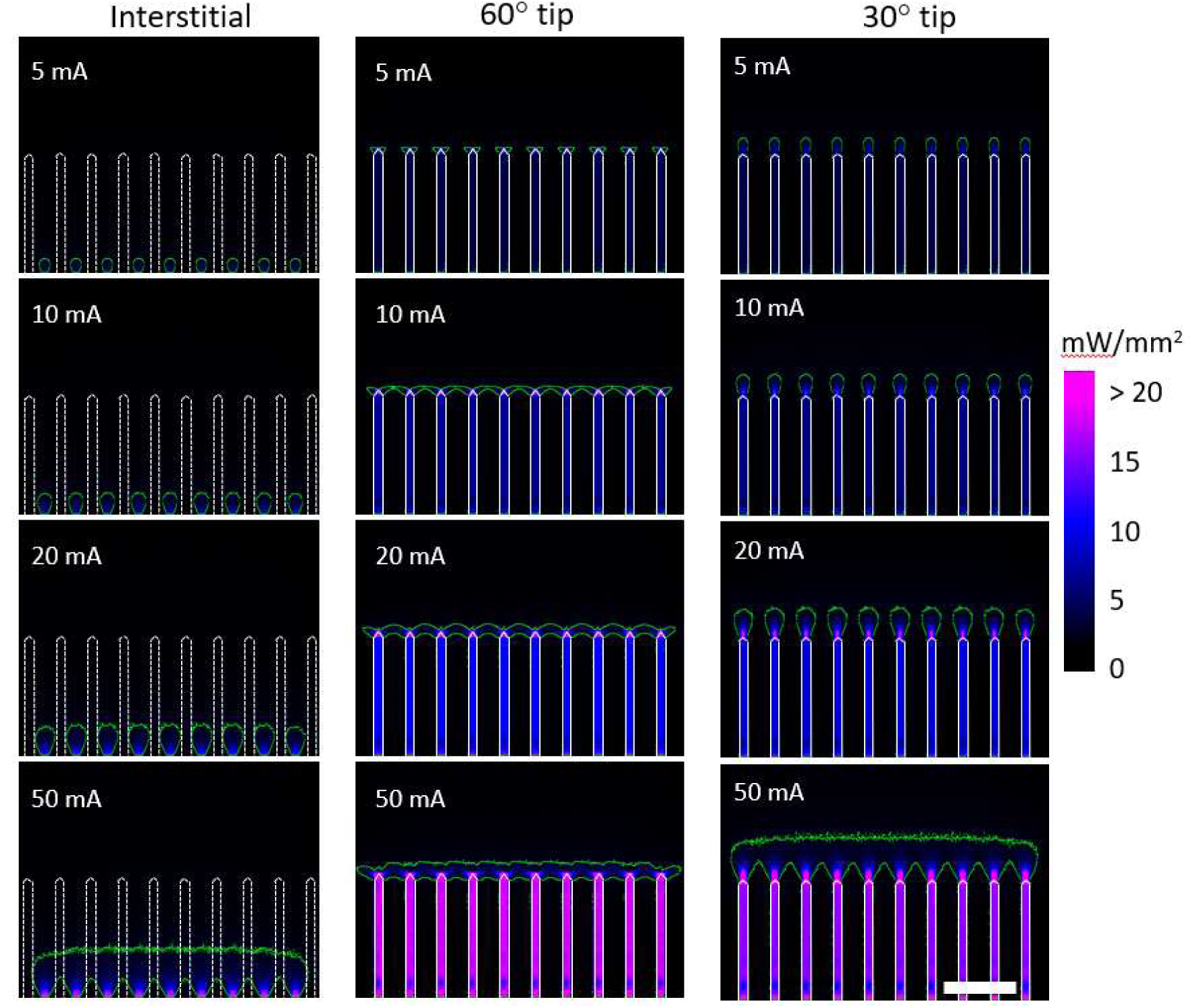
Left, 9 interstitial sites in a row simultaneously switched on. Centre, 10 needle sites with a tip of 60° simultaneously illuminated. Right, 10 needle sites with a tip of 30° simultaneously illuminated. The green line represents a 1 mW/mm^2^ threshold. The scale bar is 1 mm.

## Appendix 3

If equal irradiance is required from each needle tip the power that each µLED is driven at will differ. Figure A3 shows the power, current and voltage required for each individual µLED to output 3 mW/mm^2^.

**Figure A3:**
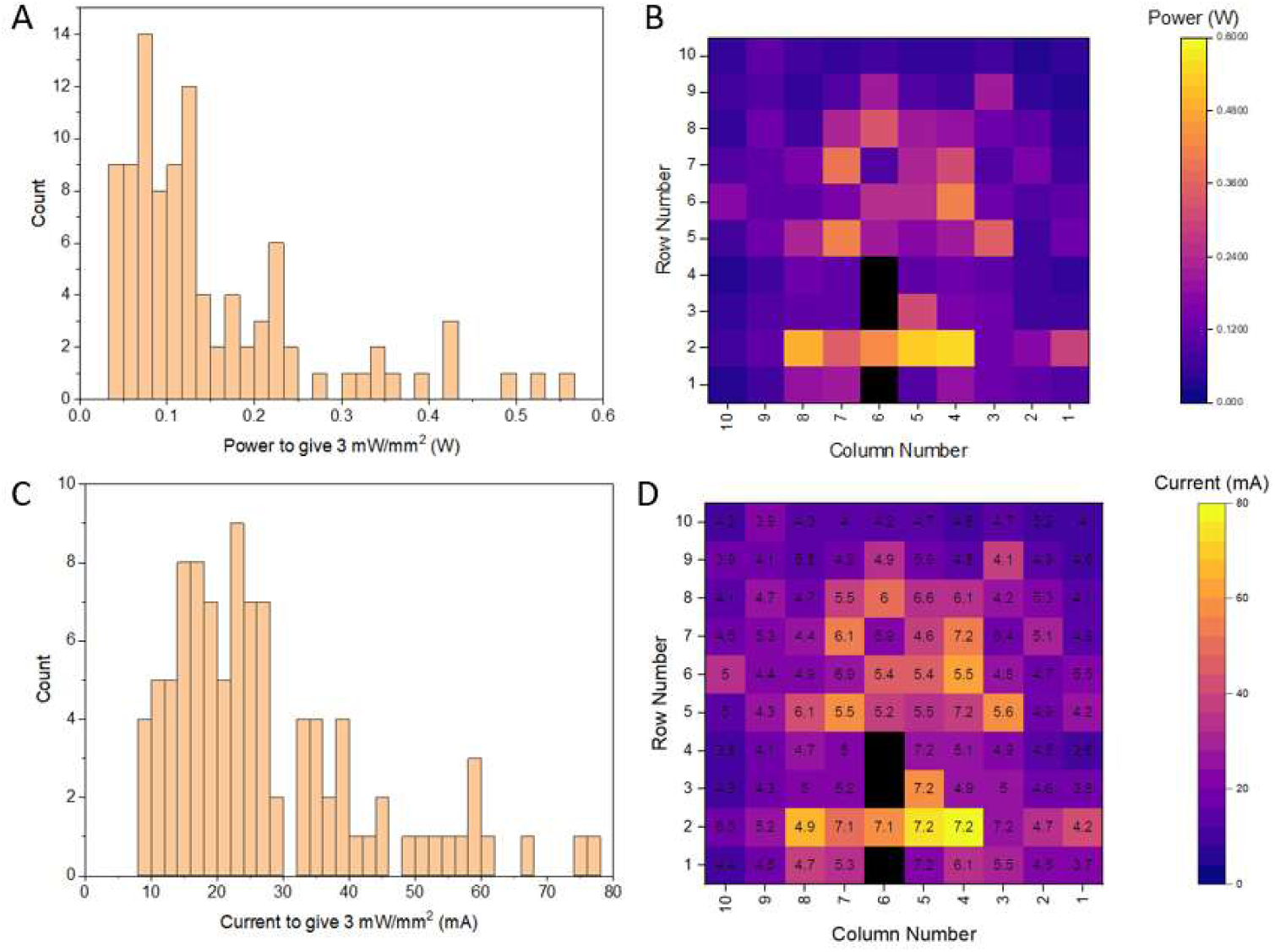
**A** Histogram of the power required to give an irradiance at the tip of the needles of 3 mW/mm^2^. **B** Map of the power required to give an irradiance at the tip of the needles of 3 mW/mm^2^. **C** Histogram of the current required to give an irradiance at the tip of the needles of 3 mW/mm^2^. **D** Map of the current required to give an irradiance at the tip of the needles of 3 mW/mm^2^. The inset number is the voltage required to drive the required current.

## Appendix 4

The thermal model was verified by comparing the measured temperature of the surface of the Dura-Gel coating directly above the µLED with the modelled temperature at the same location. The two temperature increases show good agreement for the four cases that were compared (Figure A4). Note that this is not the temperature rise in the brain, which is much smaller and quantified in Fig 4. The modelled Dura-Gel thickness can have a significant effect on the temperature at the surface of the device (the point where the thermal camera measures). In the models presented here, the total Dura-Gel thickness was taken as 1 mm (500 µm under the device, 230 µm device thickness and 270 µm on top of the device). This value was not accurately measured but estimated by analysing images after the *in vivo* study was completed.

**Figure A4:**
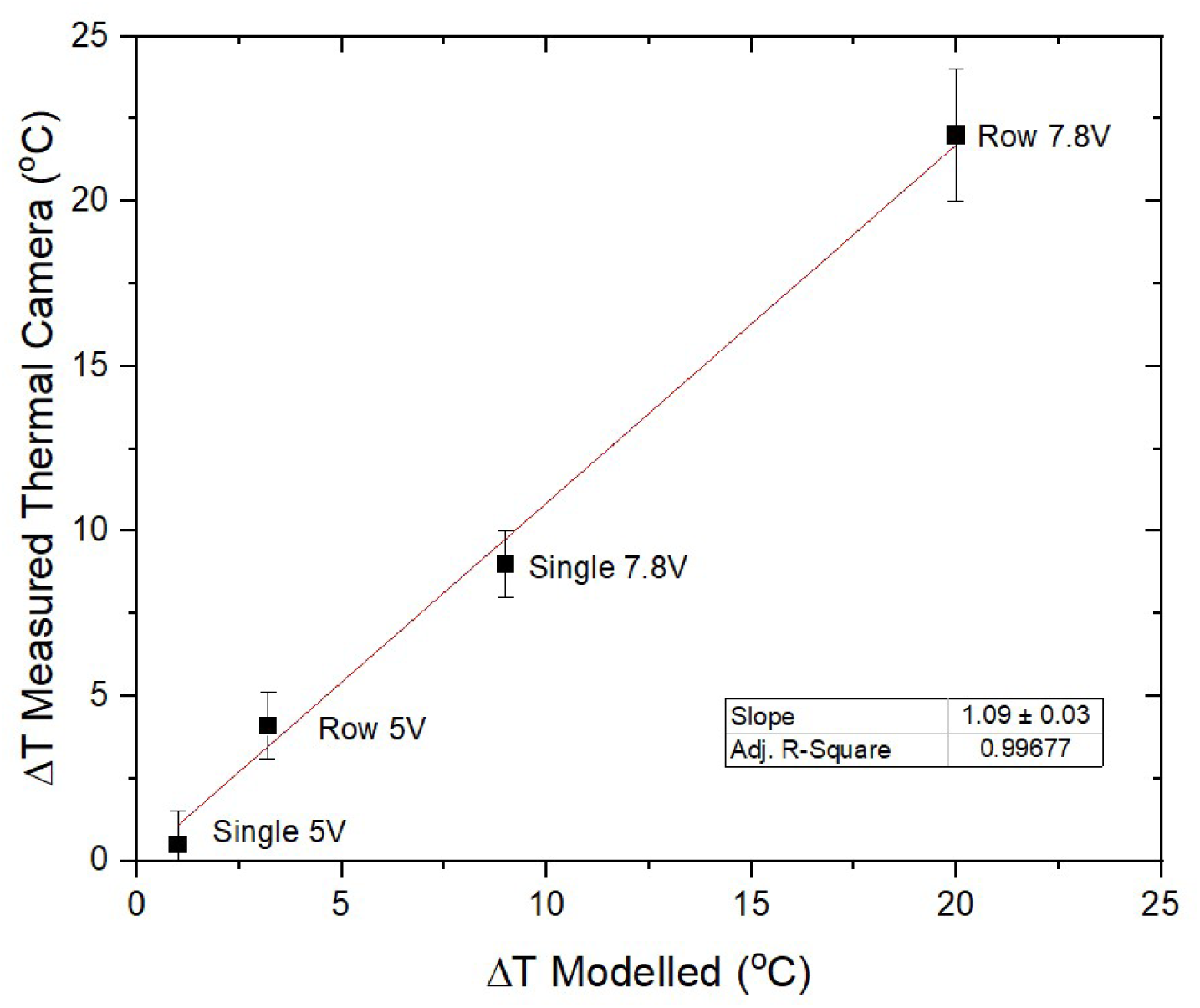
The modelled temperature at the top of the Dura-Gel surface compared to the measured temperature at the same location.

## Appendix 5

A statistical analysis of the difference between the 3 different clusters identified in the principal component analysis is shown in Figure A5 and Tables A1 and A2.

**Figure A5:**
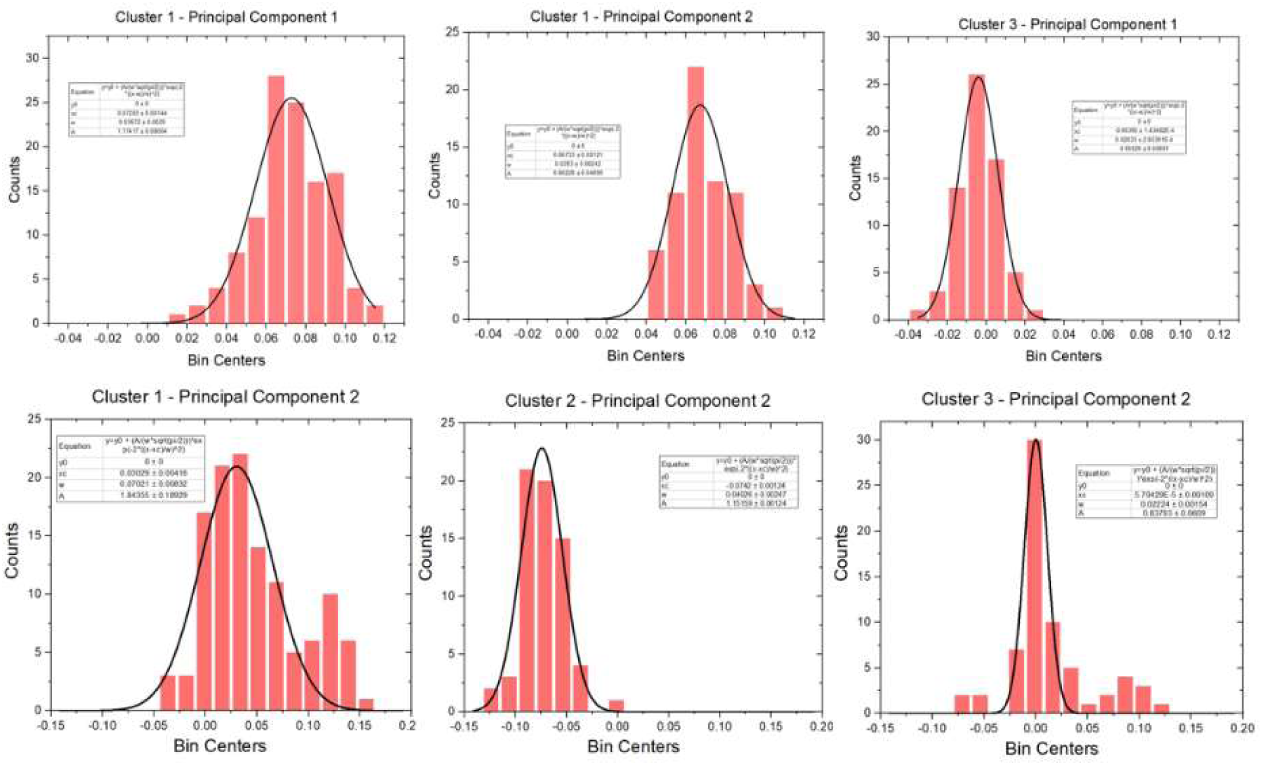
Histograms of the first two principal components for each of the clusters identified. The black line is a fit of a Gaussian function to the data. Inset: Fitting parameters for each Gaussian function.

**Table A1:** Peak position and FWHM for each of the Gaussian fits.

| Principal Component 1 |  |  |
| --- | --- | --- |
|  | Peak Position | FWHM |
| Cluster 1 | $0.073 \pm 0.001$ | $0.037 \pm 0.003$ |
| Cluster 2 | $0.067 \pm 0.001$ | $0.028 \pm 0.002$ |
| Cluster 3 | $0.004 \pm 0.0001$ | $0.02 \pm 0.0003$ |
| Principal Component 2 |  |  |
| Cluster 1 | $0.03 \pm 0.004$ | $0.07 \pm 0.006$ |
| Cluster 2 | $-0.074 \pm 0.001$ | $0.04 \pm 0.002$ |
| Cluster 3 | $0.000 \pm 0.001$ | $0.022 \pm 0.0002$ |

**Table A2:** Number of sigmas between each peak in the Gaussian fit. In this case, the sigma value was taken as the average of the sigma from each of the two Gaussian fits. This highlights that there is a large statistical difference between each of the 3 clusters identified.

| Principal Component 1 - Sigma |  |  |  |
| --- | --- | --- | --- |
|  | Cluster 1 | Cluster 2 | Cluster 3 |
| Cluster 1 |  | 0.5 | 5.9 |
| Cluster 2 |  |  | 6.1 |
| Cluster 3 |  |  |  |
| Principal Component 2 - Sigma |  |  |  |
|  | Cluster 1 | Cluster 2 | Cluster 3 |
| Cluster 1 |  | 4.2 | 1.4 |
| Cluster 2 |  |  | 5.7 |
| Cluster 3 |  |  |  |

## References

1. Ryan, T.J., et al., Memory. Engram cells retain memory under retrograde amnesia. Science, 2015. 348(6238): p. 1007–13.

2. Sahel, J.-A., et al., Partial recovery of visual function in a blind patient after optogenetic therapy. Nature Medicine, 2021. 27(7): p. 1223–1229.

3. Emiliani, V., et al., Optogenetics for light control of biological systems. Nature Reviews Methods Primers, 2022. 2(1): p. 55.

4. Jiang, S., et al., Shedding light on neurons: optical approaches for neuromodulation. National Science Review, 2022. 9(10).

5. El-Shamayleh, Y. and G.D. Horwitz, Primate optogenetics: Progress and prognosis. Proceedings of the National Academy of Sciences, 2019. 116(52): p. 26195–26203.

6. Tremblay, S., et al., An Open Resource for Non-human Primate Optogenetics. Neuron, 2020. 108(6): p. 1075–1090.e6.

7. Diester, I., et al., An optogenetic toolbox designed for primates. Nature Neuroscience, 2011. 14(3): p. 387–397.

8. Merlin, S. and T. Vidyasagar, Optogenetics in primate cortical networks. Frontiers in Neuroanatomy, 2023. 17.

9. Galvan, A., et al., Nonhuman Primate Optogenetics: Recent Advances and Future Directions. The Journal of Neuroscience, 2017. 37(45): p. 10894–10903.

10. Acker, L.C., et al., Large Volume, Behaviorally-relevant Illumination for Optogenetics in Non-human Primates. J Vis Exp, 2017(128).

11. Senova, S., et al., Experimental assessment of the safety and potential efficacy of high irradiance photostimulation of brain tissues. Sci Rep, 2017. 7: p. 43997.

12. Kim, T.I., et al., Injectable, cellular-scale optoelectronics with applications for wireless optogenetics. Science, 2013. 340(6129): p. 211–6.

13. Klein, E., et al., High-Density μLED-Based Optical Cochlear Implant With Improved Thermomechanical Behavior. Frontiers in Neuroscience, 2018. 12.

14. Scharf, R., et al., Depth-specific optogenetic control in vivo with a scalable, high-density μLED neural probe. Scientific Reports, 2016. 6(1): p. 28381.

15. Wu, F., et al., Monolithically Integrated μLEDs on Silicon Neural Probes for High-Resolution Optogenetic Studies in Behaving Animals. Neuron, 2015. 88(6): p. 1136–1148.

16. Hoffman, L., et al., High-density optrode-electrode neural probe using SixNy photonics for in vivo optogenetics. 2015 IEEE International Electron Devices Meeting (IEDM), 2015: p. 29.5.1–29.5.4.

17. Sacher, W.D., et al., Visible-light silicon nitride waveguide devices and implantable neurophotonic probes on thinned 200 mm silicon wafers. Optics Express, 2019. 27(26): p. 37400–37418.

18. Zaraza, D., et al., Head-mounted optical imaging and optogenetic stimulation system for use in behaving primates. Cell Reports Methods, 2022. 2(12): p. 100351.

19. Ruiz, O., et al., Optogenetics through windows on the brain in the nonhuman primate. Journal of Neurophysiology, 2013. 110(6): p. 1455–1467.

20. Yazdan-Shahmorad, A., et al., A Large-Scale Interface for Optogenetic Stimulation and Recording in Nonhuman Primates. Neuron, 2016. 89(5): p. 927–39.

21. Nurminen, L., et al., Top-down feedback controls spatial summation and response amplitude in primate visual cortex. Nature Communications, 2018. 9(1): p. 2281.

22. Rajalingham, R., et al., Chronically implantable LED arrays for behavioral optogenetics in primates. Nat Methods, 2021. 18(9): p. 1112–1116.

23. Abaya, T.V.F., et al., A 3D glass optrode array for optical neural stimulation. Biomedical optics express, 2012. 3(12): p. 3087–3104.

24. Lee, J., et al., Transparent intracortical microprobe array for simultaneous spatiotemporal optical stimulation and multichannel electrical recording. Nature Methods, 2015. 12(12): p. 1157–1162.

25. Afraz, A., Behavioral optogenetics in nonhuman primates; a psychological perspective. Current Research in Neurobiology, 2023. 5: p. 100101.

26. Eriksson, D., et al., Multichannel optogenetics combined with laminar recordings for ultra-controlled neuronal interrogation. Nature Communications, 2022. 13(1): p. 985.

27. McAlinden, N., et al., Multisite microLED optrode array for neural interfacing. Neurophotonics, 2019. 6(3): p. 035010.

28. Acharya, A.R., et al., In vivo blue light illumination for optogenetic inhibition: effect on local temperature and excitability of the rat hippocampus. Journal of Neural Engineering, 2021. 18(6): p. 066038.

29. Owen, S.F., M.H. Liu, and A.C. Kreitzer, Thermal constraints on in vivo optogenetic manipulations. Nature Neuroscience, 2019. 22(7): p. 1061–1065.

30. Clark, A.M., et al., An Optrode Array for Spatiotemporally Precise Large-Scale Optogenetic Stimulation of Deep Cortical Layers in Non-human Primates. bioRxiv, 2022: p. 2022.02.09.479779.

31. Yaroslavsky, A.N., et al., Optical properties of selected native and coagulated human brain tissues in vitro in the visible and near infrared spectral range. Physics in Medicine & Biology, 2002. 47(12): p. 2059.

32. Binzoni, T., et al., The use of the Henyey–Greenstein phase function in Monte Carlo simulations in biomedical optics. Physics in Medicine & Biology, 2006. 51(17): p. N313.

33. Lassen, N.A., Normal average value of cerebral blood flow in younger adults is 50 ml/100 g/min. J Cereb Blood Flow Metab, 1985. 5(3): p. 347–9.

34. Jackson, N. and J. Muthuswamy, Artificial dural sealant that allows multiple penetrations of implantable brain probes. J Neurosci Methods, 2008. 171(1): p. 147–52.

35. Potworowski, J., et al., Kernel current source density method. Neural Comput, 2012. 24(2): p. 541–75.

36. Obaid, A., et al., Ultra-sensitive measurement of brain penetration mechanics and blood vessel rupture with microscale probes. bioRxiv, 2020: p. 2020.09.21.306498.

37. Wessling, N.K., et al., Fabrication and transfer printing based integration of free-standing GaN membrane micro-lenses onto semiconductor chips. Optical Materials Express, 2022. 12(12): p. 4606–4618.

38. Gittard, S.D., et al., Two-photon polymerization of microneedles for transdermal drug delivery. Expert Opin Drug Deliv, 2010. 7(4): p. 513–33.

